# Systematic domain-based aggregation of protein structures highlights DNA-, RNA-, and other ligand-binding positions

**DOI:** 10.1101/394494

**Authors:** Shilpa Nadimpalli Kobren, Mona Singh

## Abstract

Domains are fundamental subunits of proteins, and while they play major roles in facilitating protein–DNA, protein–RNA and other protein–ligand interactions, a systematic assessment of their various interaction modes is still lacking. A comprehensive resource identifying positions within domains that tend to interact with nucleic acids, small molecules and other ligands would expand our knowledge of domain functionality as well as aid in detecting ligand-binding sites within structurally uncharacterized proteins. Here we introduce an approach to identify per-domain-position interaction “propensities” by aggregating protein co-complex structures by domain and ascertaining how frequently residues mapping to each domain position interact with ligands. We perform this domain-based analysis on ∼82,000 co-complex structures, and infer positions involved in binding DNA, RNA, peptides, ions, or small molecules across 4,120 domains, which we refer to collectively as the InteracDome. Cross-validation testing reveals that ligand-binding positions for 1,327 domains can be confidently modeled and used to identify residues facilitating interactions in ∼60–69% of human genes. Our resource of domain-inferred ligand-binding sites should be a great aid in understanding disease etiology: whereas these sites are enriched in Mendelian-associated and cancer somatic mutations, they are depleted in polymorphisms observed across healthy populations. The InteracDome is available at http://interacdome.princeton.edu.

## INTRODUCTION

The rate at which new genomes are sequenced has long since outpaced our ability to experimentally characterize the biological functions of the encoded genes and their protein products. Leveraging the fact that similar protein sequences or subsequences tend to share similar functions, computational approaches have been developed to mitigate this sequence-to-function discrepancy by rapidly detecting and modeling the sequence similarity between proteins [1]. Such homology-driven analyses of large-scale protein sequence databases have revealed many thousands of recurrent, probabilistically-modelable protein subsequences called “domains” [2, 3, 4]. These sequence-derived domains correspond to evolutionarily and functionally related substructures of proteins and are found in various modular combinations within proteins from species across the tree of life [5].

Individual protein domains are associated with specific functionalities, among the most important of which are mediating the interactions proteins make with nucleic acids, other proteins, and various other molecules in the cell. Indeed, protein–DNA and protein–RNA interactions have been found to occur via domain interfaces so frequently that factors associated with transcriptional and post-transcriptional activity are regularly classified according to their incorporation of particular nucleic acid-binding domains [6, 7]. Moreover, a significant proportion of protein–protein interactions in signaling pathways are mediated by modular binding domains [8].

Although simply knowing which domains mediate various ligand interactions has already accelerated our ability to annotate protein functions [9], pinpointing the ligand-contacting positions *within* these domains would enable a more precise analysis of the many thousands of sequenced proteins across species that contain domain instances but lack further biological characterizations. Indeed, more comprehensive knowledge of protein interaction interfaces will have considerable implications for investigating the evolution and natural variation of interaction network connectivities [10], for determining the mechanistic impact of coding variants [11], for prioritizing germline and somatic perturbations to uncover disease etiology [12, 13], and for designing targeted therapeutic drugs [14].

Identifying positions within domains that interact with ligands from sequence alone is nontrivial, as a minority of positions within a domain may be involved with ligand binding, and these positions may not be proximal with respect to the linear protein sequence (e.g., of 264 positions in the tyrosine kinase domain, only 16 noncontiguous positions contact ATP) [15]. Further, while some ligand-binding positions are largely invariant across domain instances—for example, the zinc-contacting positions in the Cys_2_-His_2_ zinc finger (C2H2-ZF) domain are required for proper domain folding and thus are highly conserved—other binding positions are not: amino acids within DNA-contacting positions in these same C2H2-ZF domains, for example, vary dramatically across domain instances to confer diverse binding specificities, and thus cannot be identified by conservation-based analyses [16].

On the other hand, analyses of three-dimensional structures of proteins co-complexed with ligands are highly accurate in identifying positions comprising interaction interfaces. Previously, co-complex structures of a single or a few manually-selected domain instances have been used as models to distinguish domain positions involved in ligand binding from those that are not [17, 18]. However, binary classifications of domain binding positions determined from single structures are not always generalizable; indeed, analyses of structurally distinct instances of some domain families have revealed that the positions involved in binding peptides or other domains can vary [19]. As such, although various databases have associated domain families with corresponding structures and bound ligands, particularly in the context of domain–domain interactions, they have largely avoided attempts to systematically determine, across multiple ligand types, the *positions* within these domains that mediate interactions [20, 21, 22, 23, 19].

Here we introduce a robust, large-scale structural aggregation approach to systematically identify positions within domains that are likely to interact with ligands. Our main contributions are as follows. First, we analyze over 82,000 protein–ligand co-complex structures in the context of domains and develop a proximity-based scoring function that determines real-valued ligand-binding propensities across individual positions in 4,120 domains; we compute per-position binding propensities separately for DNA, RNA, peptide, ion, metabolite, and other small molecule ligands. Second, we show via cross-validation testing that the resultant per-domain-position binding propensities can accurately reveal positions that bind ligands in held-out structures. Third, we utilize these **Interac**tion **Dom**ains, which we refer to collectively as the **InteracDome**, to infer interaction sites across ∼60% of human genes with high confidence, and up to ∼69% of human genes more broadly; this represents the most comprehensive resource of this type to date. Fourth, we uncover that these domain-inferred interaction sites across human proteins exhibit significant functional constraints: they are depleted for natural variants across healthy human populations, while they are enriched for Mendelian disease-associated and cancer somatic mutations. Finally, we conclude with a discussion of how our InteracDome resource can be leveraged to provide valuable, medically-relevant insights by detecting and interpreting the mechanistic effects of disease-associated coding mutations.

## MATERIALS AND METHODS

**Overview.** In this section, we describe our framework for systematically evaluating how different positions within domains are involved in mediating various ligand interactions. Briefly, we first obtain from BioLiP [24] a comprehensive collection of structures of proteins co-complexed with various ligands (Figure 1a). Each of these structures contains the three-dimensional locations of all atoms within a protein chain and all atoms within a ligand; the protein chains are also represented linearly as sequences of amino acids. We then use probabilistic sequence matching to find instances of protein domains within these protein sequences (Figure 1b). For those domain families with instances across multiple protein sequences, we aggregate their instances into a multiple sequence alignment such that each column of the alignment corresponds to a “core” domain position (i.e., a match state in the corresponding HMM profile [2]), and each row corresponds to a different domain instance (Figure 1c). Next, we analyze the structure corresponding to each domain instance. For each amino acid residue in that instance corresponding to a core domain position, we calculate the minimum Euclidean distance between any of the atoms in that residue’s side chain to any atoms in a ligand. This process leaves us with a distribution of minimum residue-to-ligand distances observed across structural instances for each domain position (Figure 1d). Finally, we distill each of these per-domain-position distance distributions into a single “binding propensity” that reflects that domain position’s propensity to bind a particular ligand (Figure 1e). We describe each of these steps in more detail next.

**Figure 1:**
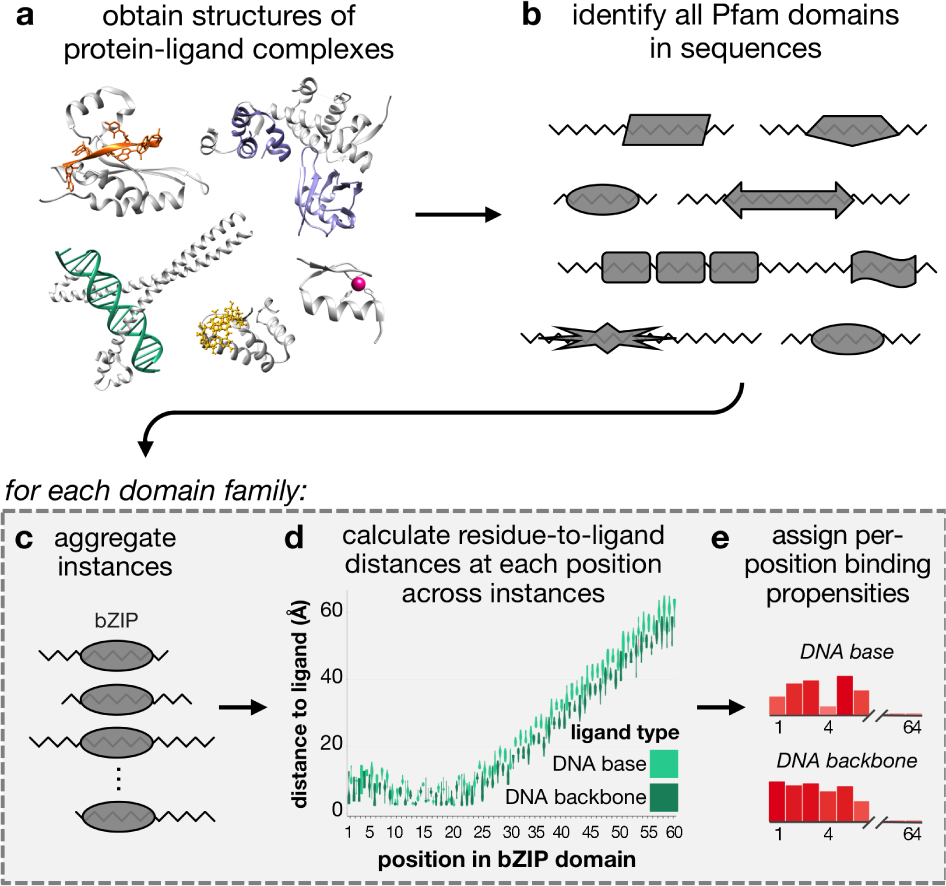
Workflow for computing per-position binding propensities for domains. **(a)** Structures of protein-ligand binding complexes are obtained from BioLiP [24]; pictured here are proteins in complex with DNA (green, PDB ID: 4auw), RNA (orange, PDB ID: 5els), peptides (purple, PDB ID: 5ibk), a zinc ion (pink, PDB ID: 1aay), and the small molecule GMP (yellow, PDB ID: 5tzd). Protein chains are colored gray. **(b)** Instances of Pfam domain families are found across BioLiP structures. For each Pfam domain family found in (b), we **(c)** aggregate all instances by ligand-binding type, **(d)** calculate distributions of minimum distances from residues to ligands, and **(e)** calculate a real-valued binding propensity for each domain position for each ligand type.

### Aggregating structural co-complexes to identify interaction domains

We downloaded crystal structures of protein–ligand complexes on June 28, 2017 from BioLiP [24]; 6,368 structures contain nucleic acid molecules, 8,314 contain peptides, and 74,726 contain additional ligands (Figure 1a). To identify domains that mediate protein interactions, we query all BioLiP protein sequences for instances of 16,712 profile Hidden Markov models from the Pfam-A database (v31.0) using HMMER (v2.3.2 and v3.1b2) [2, 25]. We restrict to domain instances that pass Pfam’s default gathering thresholds, have residues at the first and last domain positions, have the most likely residue at domain positions with information content ≥ 4 (corresponding to a distribution where the most frequent amino acid appears with ∼95% probability), and contain at least one residue annotated by BioLiP to be involved in ligand binding (Figure 1b).

### Differentiating and grouping ligand types

We classify BioLiP ligands into biologically relevant groups. Protein residues responsible for determining DNA-and RNA-binding specificity often contact nucleic acid bases, whereas residues that primarily contact the backbones of nucleic acid molecules may be more important for the stability and affinity of the binding complex [17, 26]. As BioLiP groups all DNA and RNA ligand molecules together, we reanalyze the original co-complex structures to characterize ligand atoms based on the presence (RNA) or absence (DNA) of the 2’-hydroxyl in the ribose sugar; these ligands respectively occur in 2,244 and 4,124 co-complex structures. We further group these ligand atoms into RNA base, RNA backbone, DNA base or DNA backbone.

The 205 ligands from 34,705 co-complex structures with “ion” in their full names are assigned to the ion group; the remaining 21,078 ligands from 58,650 co-complex structures are grouped as small molecules. BioLiP already excludes molecular artifacts from crystallization buffers, yet does not explicitly differentiate cognate (i.e., naturally occurring *in vivo*) from non-cognate ligands. To highlight domain positions whose residues comprise metabolically-relevant and/or potentially druggable binding pockets, we further categorize small molecules as follows. Any small molecule ligand with a Tanimoto coefficient ≥ 0.9 (Open Babel Package, v2.4.1) [27, 28] between its SMILES string (wwPDB’s Chemical Component Dictionary, v3.30) and the SMILES strings of one of 29,332 endogenous human metabolites (Human Metabolome Database, v3.6) [29] and/or 7,162 drugs (DrugBank, v5.0.1) [30] is respectively classified as metabolite and/or druglike; these ligand types respectively occur in 19,117 and 43,535 co-complex structures.

### Computing proximity-based positional binding propensities

For each domain with corresponding co-complex structure(s) (which we refer to as the *modelable* set of domain–ligand interactions), we assess the ligand-binding propensities of individual domain positions by aggregating protein–ligand atom proximity information across their corresponding co-complex structures (Figure 1c-e). For each instance of the domain in a BioLiP structure that contains at least one residue in contact with a particular ligand type, for each of its residues corresponding to a Pfam domain match state, we compute the minimum Euclidean distance between heavy (i.e., non-hydrogen) side-chain atoms to any heavy ligand atom. We aggregate these distances by domain position across all BioLiP instances for the same domain–ligand type pair, resulting in a per-domain-position distribution of minimum distances to the ligand (Figure 1c-d). We next determine what fraction of these distances are within 3.6Å of the ligand (Figure 1e); however, to proportionally minimize the contribution of structural instances with highly redundant sequences, we apply the Henikoff and Henikoff sequence weighting scheme [31] to the domain instances. Thus, intuitively, our per-domain-position binding propensity computes the (weighted) fraction of times a residue in that position is within 3.6Å of the ligand type considered. Note that the distance of 3.6Å will capture both hydrogen bonds (2.6-3.3Å) and van der Waals contacts (2.8-4.1Å), but will eliminate water-mediated interactions (as in [32]). We find that residue-to-ligand proximity cutoffs between 2.5 and 6.0Å do not significantly alter downstream results.

### Cross-validation testing of domain-to-ligand distance consistencies and positional binding propensities

We refer to each domain–ligand type pair with at least three distinct (i.e., non-redundant) sequences across separate PDB entries as belonging to the *modelable-NR* set. For each domain–ligand pair in the modelable-NR set, we evaluate both the consistency of its structural interface as well as the accuracy of our aggregation approach in identifying ligand-binding positions in cross-validation.

First, to evaluate consistencies of domain–ligand structural interfaces, for each domain–ligand pair in the modelable-NR set, we first randomly split its structural instances into two folds. We next compute, across all instances within each fold, the average minimum residue-to-ligand distance at each domain position. Finally, we compute the Pearson’s correlation coefficient (PCC) between the two resulting domain-length vectors. We report the average PCC achieved across ten repetitions of this process as the consistency of the domain–ligand structural interface.

Second, to test the predictive power of our approach, for each domain–ligand type pair in the modelable-NR set, we randomly divide its structural instances into up to ten folds. For each domain instance *i* in each hold-out fold in turn, we examine the structure to assign a binary vector where 1s and 0s respectively indicate domain positions whose residues are or are not in contact with the ligand (i.e., as annotated by BioLiP). Binding propensities are then calculated as before from instances in the remaining folds. For each position in each instance in the hold-out fold, we use the corresponding positional binding propensity, computed from the other folds, as its “score.” We then rank in descending order all positions within the hold-out fold by score, with higher ranking positions corresponding to the more confident predictions of binding. As we iteratively decrease the score threshold used to predict whether a position is binding, we compute precision and recall with respect to the known binding (true positive) and non-binding (true negative) positions inferred from the actual structures in the held-out set. This allows us to compute a precision-recall curve (PRC) for each domain–ligand interaction. We refer to the set of domain–ligand interactions that achieved a cross-validated precision of at least 0.5 at some threshold as the *confidently modeled* set. We also compute the area under under the PRC (AUPRC), and compare it to an average baseline AUPRC corresponding to the fraction of binding positions in the held-out set.

We note that instances of the same domain family have by definition clearly identifiable sequence similarity and thus can have highly similar amino acid sequences. Nevertheless, as an alternate cross-validation test, for each domain–ligand type pair in the modelable set, we also try to divide all its instances within BioLiP into groups such that the amino acid sequence identity between instances in different groups is <90%. We repeat the steps above to determine domain–ligand structural consistencies and cross-validated precisions and recalls by dividing these groups of BioLiP instances with sequence identity ≥90%—rather than individual instances—into folds as before; these results are reported in the Supporting Information section.

### Human protein, natural variation and disease mutation datasets

Protein sequences, corresponding cDNA sequences and corresponding genomic coordinates for 104,295 known and predicted human protein isoforms encoded by 23,043 genes were downloaded from Ensembl (build GRCh38.p10). We consider the subset of 89,024 protein isoforms from 22,712 human genes where the genomic DNA sequence matched the cDNA sequence, the cDNA sequence translated to the protein sequence with ≤5% sequence mismatch, and the protein transcript was not annotated with ‘decay’ nor ‘pseudogene.’ We functionally classify single nucleotide variants (SNVs) with respect to the longest protein isoform for each gene by mapping SNVs onto Ensembl cDNA sequences and translating to proteins.

Naturally occurring exonic SNVs from 123,136 healthy humans were downloaded from the Genome Aggregation database (gnomAD, v2.0.2) [33]. We restrict to the set of 194,868 common missense SNVs that are found with frequency ≥ 0.001 across any gnomAD subpopulation. We also obtained 28,242 missense germline disease mutations affecting 24,823 sites across the canonical protein isoforms of 2,590 human genes from UniProtKB’s Humsavar database (v2017_04) [34] and augmented this set with an additional 1,912 validated missense germline disease mutations occurring an additional 159 human genes from OMIM (v2011_02) [35].

We also downloaded all open-access TCGA somatic SNV data and RNA-seq expression data from NCI’s Genomic Data Commons on July 15, 2017 [36, 37]. We exclude all SNVs occurring after a frameshift or nonsense mutation in the corresponding tumor sample and all SNVs from genes that were expressed at <0.1 TPM (in the corresponding tumor sample or on average across other tumor samples of the same tissue type when expression data was missing). These steps resulted in a filtered set of 1,171,890 missense somatic SNVs across 18,627 genes using data from 10,037 tumors across 33 cancer types. Finally, 1,209 known cancer driver missense SNVs were downloaded from the Database of Curated Mutations (DoCM, v3.2) [38].

### Inferring putative ligand-binding positions in human proteins

We query the longest protein isoform of each human gene and infer ligand-binding positions in these proteins in three ways. First, we extract human proteins from BioLiP, and obtain the residues identified in this database to interact with ligands. Next, we transfer structural binding information from BioLiP to human proteins with high sequence similarity, as described previously [13]. Finally, for each domain where we have estimated per-position binding propensities, we find matches to this domain in human sequences using HMMER as described above, and transfer the ligand-binding propensities to any protein residue that corresponds to a core domain position. In practice for this last step, only confidently modeled domain–ligand pairs are used. For each of these domain–ligand pairs, the threshold to define ligand-binding positions is chosen as the value that resulted in cross-validated precision ≥0.5, as described above.

### Determining significance of overlap with inferred ligand-binding sites

Given a set of sites of interest in human proteins (e.g., sites harboring common missense SNVs across populations) and a set of putative ligand-binding sites in human proteins, we determine whether the overlap between these two sets is significantly larger or smaller than what is expected by chance alone using the Poisson binomial distribution. Here, the *N* sites of interest across proteins are modeled as *N* independent Bernoulli trials, where the *p*_1_, …, *p_N_* probabilities of “success” (i.e., overlap with the putative binding sites) for each trial are non-uniform. Specifically, each success probability *p_i_* is equal to the proportion of putative binding sites in the protein where the site of interest *i* occurs; this way, sites of interest that occur in proteins with a large proportion of putative ligand-binding sites will not bias global trends. We determine if *K* —the number of sites of interest observed to overlap with the putative binding sites—is significantly greater than or less than we would expect by chance by respectively computing Pr(*X* ≥ *K*) and Pr(*X* ≤ *K*) using the Poisson binomial implemented in R’s poibin package [39]. *P*-values computed as 0 are reported as 1e-15, the lowest non-zero *p*-value we achieved using poibin.

## RESULTS

Our fully automated procedure to build the InteracDome resource identifies 4,120 domain families with modelable ligand interactions with one or more instances across structural co-complexes. Of these, 2,055 domain families have at least three non-redundant instances in complex with the same ligand type across distinct PDB structures; these domain–ligand interactions comprise the modelable-NR set. Within the modelable set of domain interactions, 572 domain families are co-complexed with DNA, 491 are co-complexed with RNA, 2,553 are co-complexed with ions, 840 are co-complexed with other peptides, and 2,830 are co-complexed with one or more small molecules. In the modelable-NR set, 168, 189, 1,019, 225, and 1,341 domain families are respectively co-complexed with DNA, RNA, ions, peptides, and/or small molecules.

### Case studies: InteracDome includes well-known interaction domains and recapitulates known ligand-binding domain positions

We begin by ascertaining how well the interaction domains profiled in the InteracDome cover known ligand-binding domains. Towards this end, we compiled a list of 54 DNA-binding domain families from the Thornton Lab review [40], 12 RNA-binding domain families from the review by [41], and 78 human peptide-binding domain families listed on the Pawson Lab site (http://pawsonlab.mshri.on.ca). We find that our modelable set of domain–ligand interactions includes all the DNA-binding domain families, all the RNA-binding domain families, and ∼85% of postulated protein-binding domain families, many of which have been particularly difficult to structurally characterize due to the low affinity and transience of protein–protein interactions in signaling pathways [42]. Of these known ligand-binding domains, 36 (67%) DNA-binding domains, 8 (50%) RNA-binding domains and 39 (50%) peptide-binding domains are found in the modelable-NR set; these numbers are 36 (67%), 6 (50%) and 36 (46%), respectively, for known ligand-binding domains found in the confidently modeled set.

Next, we turn our attention to how well our per-domain-position binding propensities identify manually curated ligand-binding domain positions. In particular, domain positions involved in ligand binding have previously been established for a few well-studied domains using one or a few structural co-complexes. Intuitively, our method automates this approach at a much larger scale; thus, we expect that domain positions assigned high binding propensities by our method will largely be in agreement with previous knowledge of domain binding. We highlight below a few well-studied nucleic acid-, peptide-and metabolite-binding domains to show that indeed, when we compare InteracDome binding propensities with literature-curated knowledge of domain–ligand binding, high propensity domain positions recapitulate known interaction positions.

We first consider three nucleic acid binding domains. The C2H2-ZF domain is known to specify its DNA targets via four DNA-base contacting positions (−1, 2, 3 and 6 in the *α*-helix contacting DNA) [32]; these four positions have the highest DNA base-binding propensities for this domain in the InteracDome. Additionally, there are two highly conserved cysteines and histidines that coordinate the zinc ion—required for proper domain folding—and these four positions have our highest ion-binding propensities (Figure 2a, Figure S1a). In the DNA-binding homeodomain, our highest DNA base-binding propensities correspond to positions 45–46, 49–50 and 53–54 in the DNA recognition helix, followed by positions 1–4 in the N-terminal arm (Figure 2b, Figure S1b); these are known specificity-determining positions in the domain [43, 18]. In the RNA-binding pumilio domain, the highest RNA base-binding propensities are found in positions 14, 16, 17 and 20 of the repeating alpha-helix section (Figure 2c, Figure S1c). Indeed, positions 16, 17 and 20 confer RNA-binding specificity, and position 14 contacts RNA backbone ribose rings, likely affecting binding [26].

**Figure 2:**
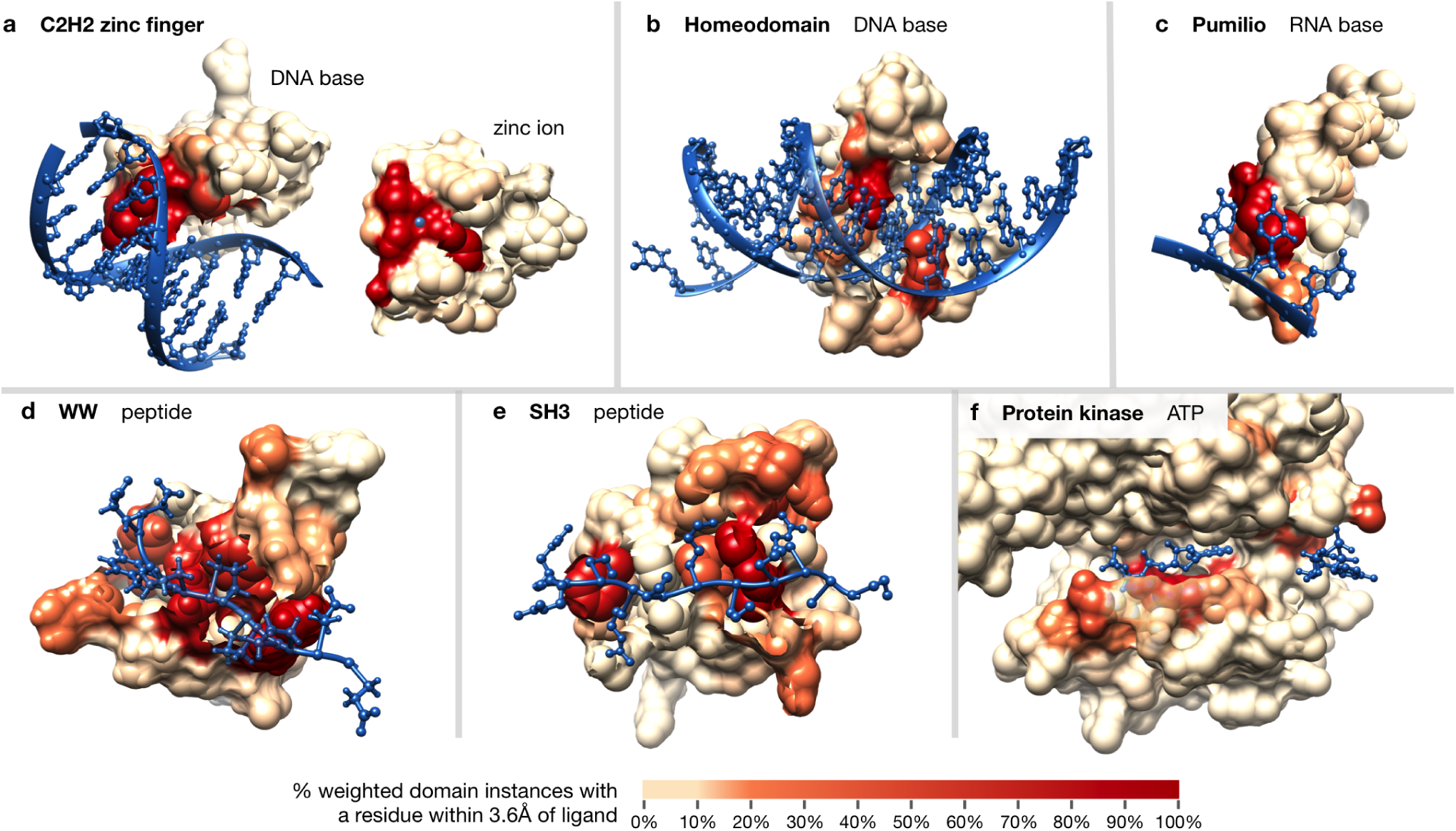
Examples of domains scored according to ligand-binding propensity. Residues within interaction domain structures are colored according to their ligand-binding propensities; domains pictured include: **(a)** C2H2-ZF domain (PF00096, PDB ID: 1aay, second domain of chain A), the zinc ion is shown on top of the domain for visibility, **(b)** Homeodomain (PF00046, PDB ID: 1ig7), **(c)** Pumilio domain (PF00806, PDB ID: 1m8w, third domain of chain B), **(d)** WW domain (PF00397, PDB ID: 2n1o), **(e)** SH3 domain (PF00018, PDB ID: 2bz8), and **(f)** protein kinase domain (PF00069, PDB ID: 1csn, subdomains I-V, both ATP molecules from two units shown).

We next examine InteracDome binding propensities for two peptide-binding domains. In the WW domain, our highest peptide-binding propensities are found at positions 17, 19, 21, 24, 26 and 28—all corresponding to known binding residues. The next highest peptide-binding propensities identify positions 8, 10 and 11—all known to confer binding specificity differences between type I and IV domains (Figure 2d, Figure S1d) [44]. Our high propensity sites for the SH3 domain are also relevant for peptide binding. In particular, our highest propensity positions are 4, 6, 32 and 47—the four most conserved peptide-binding sites—and positions 9–13, 27–30, 34 and 45—all known to be important for distinct peptide-binding specificities (Figure 2e, Figure S1e) [45].

Finally, our approach also recapitulates ATP-binding positions in the kinase domain. Our high small molecule propensities include several residues from subdomains I–VII that are responsible for interacting with and anchoring ATP’s adenine ring (i.e., positions 7, 15, 28, 77–80, 84), *α*, *β* and γ phosphates (i.e., positions 11–13, 30, 128, 141), and ribose hydroxl group (i.e., position 127) [15]. We also highly rank an additional six sites within three amino acids of a known binding position (Figure 2f, Figure S1f).

### Domain-to-ligand proximities are consistent across instances

We next show, in a systematic analysis, that structural interfaces between domains and their ligands in the modelable-NR set tend to be conserved. Briefly, we compare the residue-to-ligand distances across different structural instances of a domain–ligand interaction type to each other (see Materials and Methods). We find that analogous positions across domain instances indeed tend to have similar distances to ligands: the median PCC of domain–ligand interactions is 0.97 and the PCC ≥ 0.8 for 90% of domain–ligand interactions (Figure 3,Figure S2). Thus, domains tend to have highly consistent structural interaction interfaces with the same ligand type across instances.

**Figure 3:**
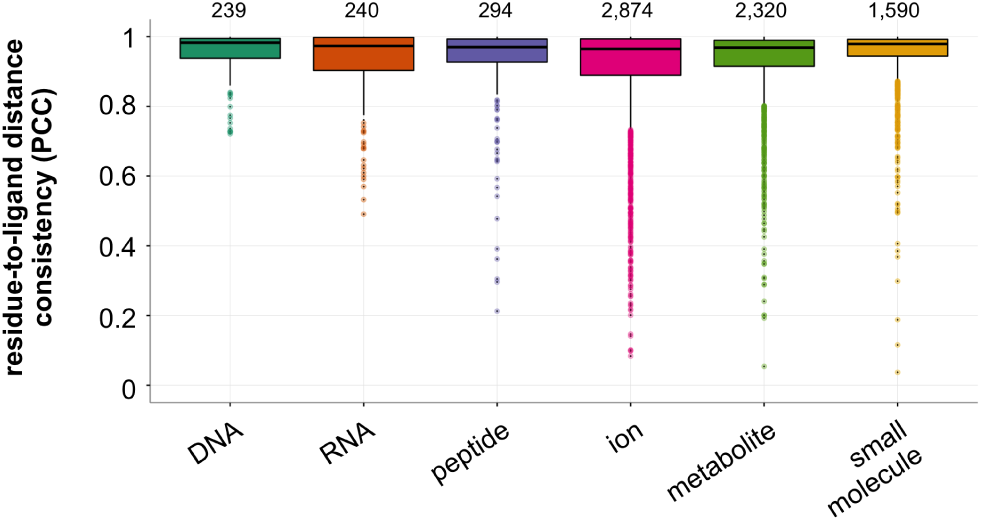
Domain-to-ligand distance consistencies. Structural instances across BioLiP for each domain–ligand type in the modelable-NR set are randomly split into two folds. Shown are Pearson’s correlation coefficients (PCCs) of the average residue-to-ligand distances across each domain position between the two folds, averaged across ten repetitions. Relative domain-to-ligand distances across domain positions tend to be highly consistent between structural instances of the same domain–ligand pair. Total number of domain–ligand interactions included in each boxplot are listed above the corresponding distributions.

As we have just shown, domains generally tend to interact with their ligands in a relatively consistent fashion. However, some interaction domains are also known to have multiple binding modes, where different combinations of domain positions confer binding specificity and affinity [44, 45]; these cases cannot be detected by examining structural domain instances in isolation, but may be revealed using our aggregation approach. To get a better idea of how much domain positions vary with respect to their roles in ligand binding, for each domain–ligand type pair in the modelable-NR set, we use empirical bootstrapping of structural instances with 1,000 repetitions to obtain standard errors of all binding propensities; smaller standard errors indicate a domain position’s consistent role in binding, whereas larger bootstrapped standard errors indicate a position’s more variable role in binding. Importantly, standard errors tend to be low for the full range of ligand-binding propensities (Figure 4a, Figure S3a). Moreover, as expected, positions with more extreme binding propensity values (i.e., ≥0.95 or ≤0.05) tend to have lower standard errors. Conversely, positions with intermediate binding propensities also exhibit more variation in their estimates in bootstrapped samples. We also show that with more sequentially-distinct structural examples of domain–ligand complexes, the standard errors of computed binding propensities decrease overall (Figure 4b, Figure S3b). This indicates that aggregating information across structural domain instances allows us to infer which positions within domains bind particular ligands more confidently than we could if we were limited to only one or a few structural examples.

**Figure 4:**
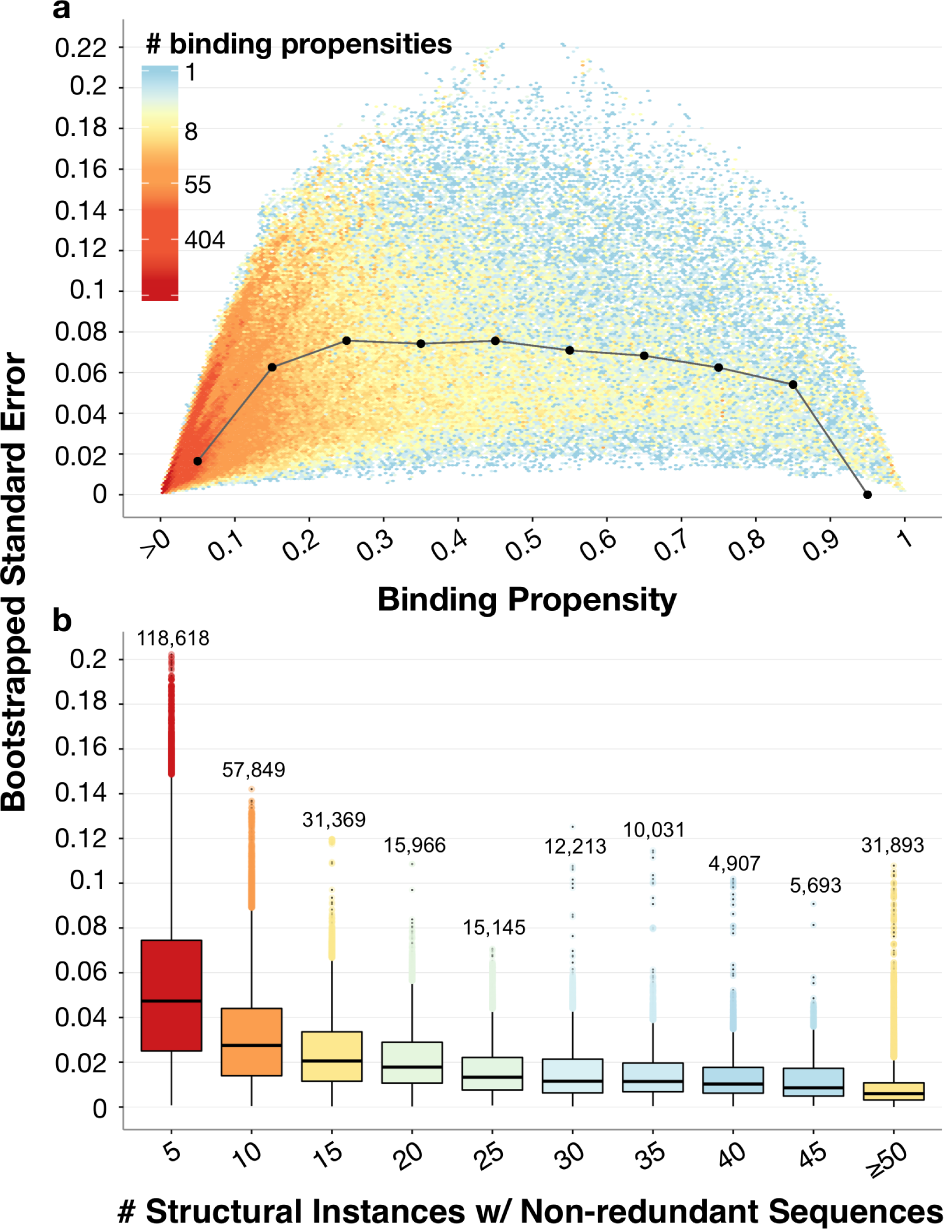
Bootstrapped standard errors of binding propensities. **(a)** For each domain position with a positive binding propensity in each domain–ligand interaction pair in the modelable-NR set, we plot its ligand-binding propensity (*x*-axis) and the standard error of this propensity (*y*-axis), computed as the standard deviation of its ligand-binding propensity as measured over 1,000 bootstrap samples. Distribution medians at each binding propensity decile are shown as black dots and are connected by gray lines for visual effect. **(b)** Bootstrapped standard errors decrease as the number of structural domain instances with non-redundant sequences increase, illustrating the ability of our structural aggregation approach to determine how domain positions are generally involved in ligand binding. Boxplots are colored according to the relative size of each distribution; the number of total domain positions, across domain–ligand type pairs, is listed above each boxplot.

### Cross-validation highlights power of binding propensities

We next evaluate how well the binding propensities for a given domain–ligand type pair indicate positions involved in binding across previously unobserved structural instances. To measure this, we employ cross-validation to compute PR curves for each domain–ligand pair in the modelable-NR set, and compare the AUPRCs to corresponding baseline AUPRCs (see Materials and Methods). We find that the average (across folds) actual AUPRCs are typically substantially higher than their corresponding baseline AUPRCs, particularly for domain–ion interactions which tend to involve far fewer domain binding positions and thus have lower baseline AUPRCs (median fold improvement of actual over baseline AUPRCs = 24.0, fold improvement ≥ 5 for 95.5% of domain–ligand interactions, Figure 5a). We also note that the cross-validated precisions for domain–ligand type pairs in the modelable-NR set tend to be high across a range of binding propensity cutoffs (Figure 5b). Moreover, when we repeat this process with stricter fold divisions, ensuring that structural instances of domain–ligand interactions in separate folds have <90% sequence similarity, we again find that the improvement of actual over baseline AUPRCs remains high (median fold improvement of actual over baseline AUPRCs = 21.9, fold improvement ≥ 5 for 93.8% of domain–ligand interactions, Figure S4). This benchmarking demonstrates that our binding propensities can be used to infer domain binding positions across previously unseen, sequentially diverse structural instances. For the remainder of our analysis, we consider a filtered *confidently modeled* set of 10,299 domain–ligand interactions, involving 1,327 distinct domains, from the modelable-NR set that achieve a cross-validated precision ≥0.5 at some binding threshold.

**Figure 5:**
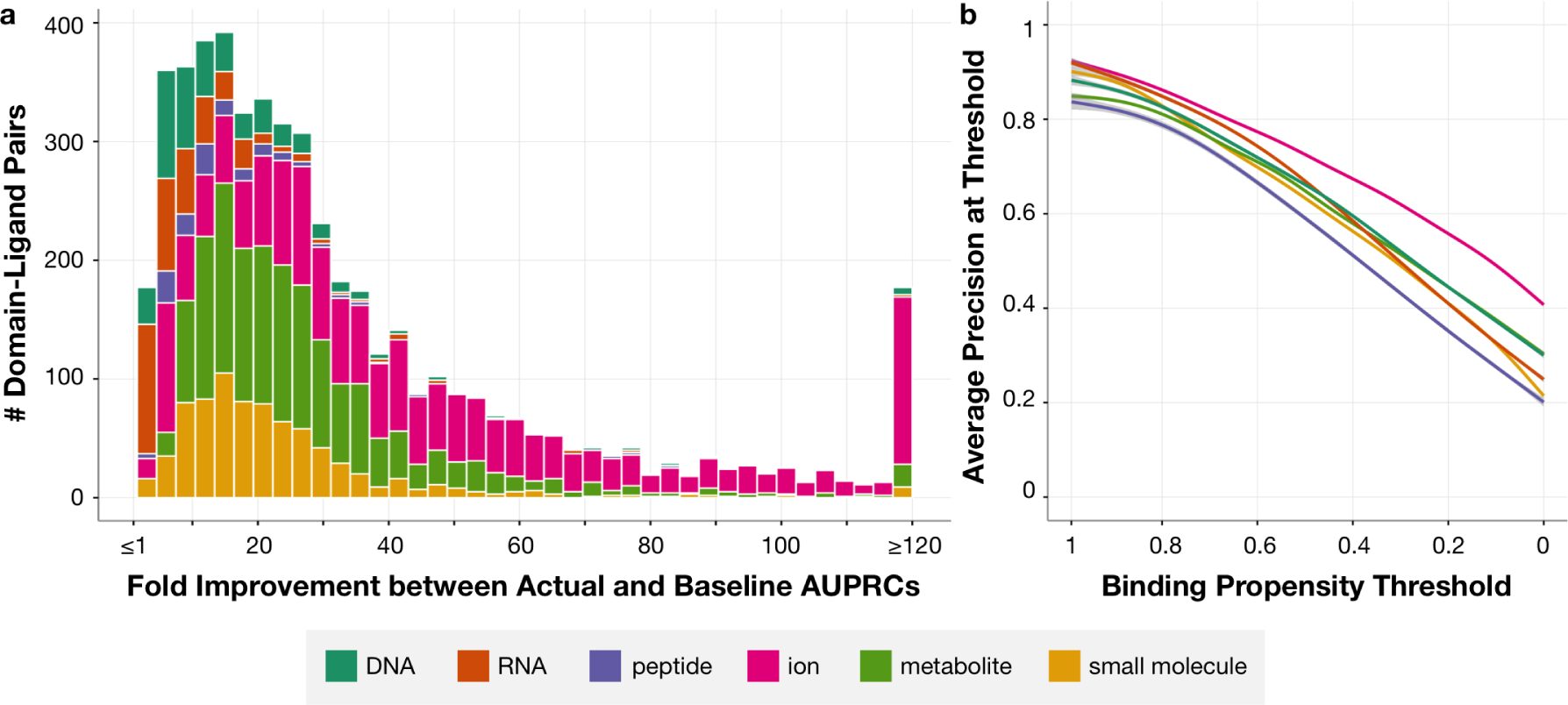
Cross-validation testing of binding propensities. Accuracy of each domain–ligand interaction from the modelable-NR set is measured as the average AUPRC in cross-validation with up to 10 folds. **(a)** For each domain–ligand pair, we compute the fold change between the actual AUPRC and a baseline AUPRC corresponding to the fraction of binding positions in the hold-out set. **(b)** We use each positional binding propensity computed across domain–ligand pairs as a threshold to distinguish predicted binding from non-binding domain positions, and we measure the precision achieved in each held-out set of domain–ligand structural instances using each of these thresholds. Shown is the average precision computed across domain–ligand interactions in the modelable-NR set at binding propensity thresholds varying from 1 (highest) to 0 (lowest).

### Analysis of InteracDome-inferred binding sites in human

We next use our InteracDome resource to infer sites in human proteins that may be involved in interactions with DNA, RNA, peptides, ions, metabolites and small molecules.

#### Domain-based approach doubles coverage of human genes with modeled interactions

As of June 2017, only 2,871 (12.6% of 22,712 total) human genes were associated with biologically relevant protein– ligand complex structures in BioLiP. Homology modeling as described in [13] allows us to infer binding residues in an additional 2,990 genes. Together, these two approaches cover 25.8% of all human genes (Figure 6a) [46, 47, 13]. We note that there are an additional 786 human genes associated with co-complex structures in the Protein Data Bank (PDB) [48], but these proteins are either complexed with non-biologically relevant ligands or with peptides longer than 30 residues (which are not included in our analysis).

**Figure 6:**
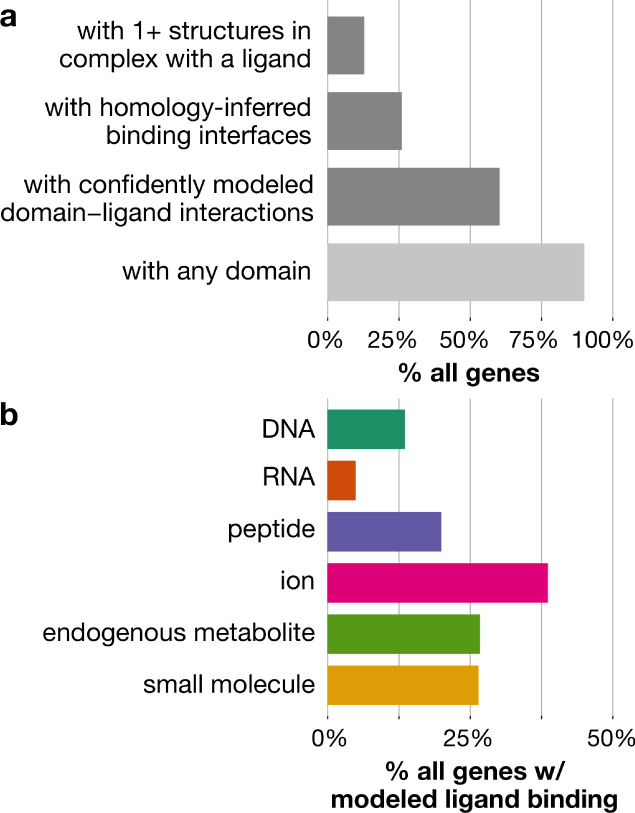
Interaction domain-based coverage of human genes. **(a)** Structural and domain-based coverage of 22,712 human genes. Dark gray bars indicate ways by which to structurally model protein interactions. **(b)** Percentages of genes estimated to interact with specific ligand types using the confidently modeled set of domain–ligand interactions.

Approximately 90% of human genes contain complete instances of ∼6,000 Pfam domain families. Of course, not all of these domains have associated structural co-complex information, and thus neither their roles in mediating binding nor which positions within them are involved in ligand binding are known. However, 13,654 (60.1%) genes contain InteracDome domain instances with confidently modeled interactions or homology-inferred binding interfaces. Including any modelable interaction domain with 10+ or 1+ instances in BioLiP, rather than only domains with confidently modeled interactions, respectively covers 61.4% and 68.7% of human genes. Our InteracDome resource thus represents a 2.3-to 2.7-fold increase in coverage over current state-of-the-art approaches to infer putative interaction sites across human genes. Furthermore, our approach covers a diverse range of interaction types, as substantial fractions of these genes have confidently modeled sites involved in binding DNA, RNA, peptide, ions, metabolites and other small molecules (Figure 6b). Altogether, the InteracDome represents a considerable improvement in our ability to infer diverse protein–ligand interaction sites across large numbers of proteins across species.

#### Putative Binding Sites are Depleted of Natural Variants, Enriched for Disease Mutations

Because the vast majority of proteins’ functions are carried out through specific interactions, even rare DNA variants or mutations that alter interaction-mediating protein residues can have critical impacts in human disease. As such, we expect inferred protein interaction sites to be relatively conserved across healthy human individuals, whereas we would expect these same sites to be perturbed across individuals with disease [13]. To determine whether our InteracDome-inferred binding residues exhibit these expected functional constraints, we perform an initial analysis on missense mutations where we consider any protein residue overlapping a domain match state with a corresponding binding propensity that resulted in a cross-validated precision ≥ 0.5 to be a confidently modeled“putative” ligand-binding site (see Materials and Methods).

We first assess whether commonly varying sites across healthy human individuals tend to globally overlap with putative ligand-binding sites as expected across the human proteome [33]. We find that the overlap between commonly varying sites and confidently modeled domain-inferred binding sites is significantly *less* than expected by random chance (*p* < 1e-15, Poisson binomial test, Figure 7a). This global trend indicates that sites identified by InteracDome as potentially ligand-binding are generally conserved across healthy individuals, in accordance with what we would expect and further demonstrating the utility of our resource to highlight functionally important protein interaction positions.

**Figure 7:**
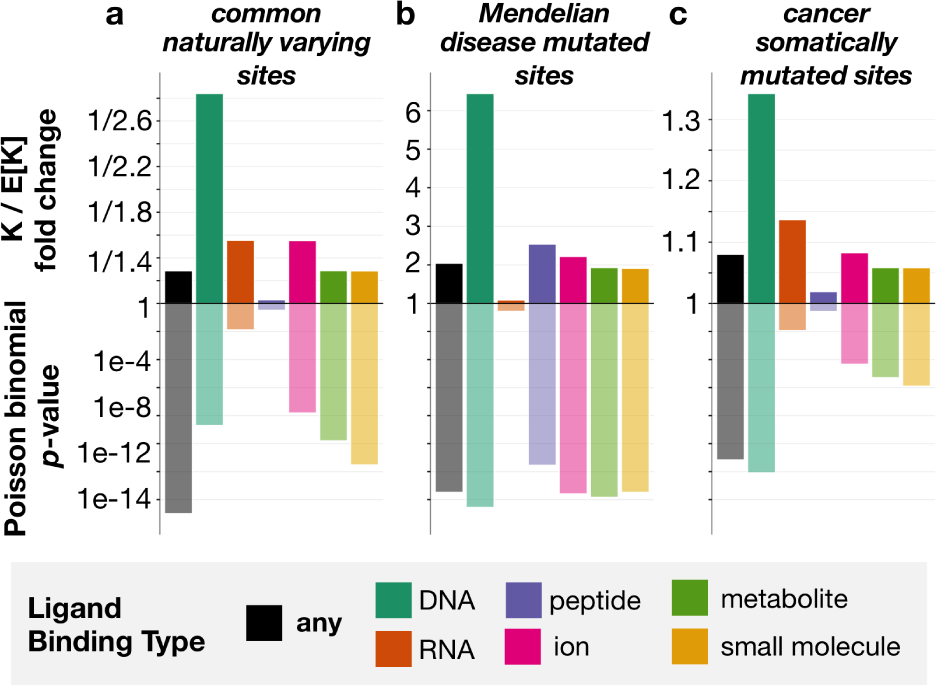
Natural variants show opposite trends from disease mutations with respect to ligand-binding sites. Putative ligand-binding sites correspond to protein positions overlapping domain match states whose binding propensities resulted in a precision of at least 0.5 in cross-validation testing (i.e., confidently modeled interactions, see Materials and Methods). Shown on the *y*-axis (top) is the fold change between the observed (K) and expected (E[K]) numbers of InteracDome-inferred putative binding sites (any type, DNA, RNA, peptide, ion, metabolite or small molecule) and other sites of interest (common naturally varying, Mendelian disease mutated, cancer somatically mutated). We compute the significance of this overlap (*y*-axis, bottom) using the Poisson binomial distribution. **(a)** Putative ligand-binding sites exhibit a significant lack of overlap with commonly varying sites across human proteins. **(b)** Conversely, putative ligand-binding sites overlap significantly with sites harboring Mendelian disease mutations. **(c)** Protein sites harboring missense cancer somatic mutations also overlap significantly with putative ligand-binding sites, suggesting that these sites are preferentially altered in human cancers.

We next consider whether protein positions harboring known disease-associated mutations overlap with these same putative binding sites. We uncover that Mendelian disease-mutated sites coincide with putative binding sites far *more* than expected by chance (*p* < 3.3e-14, Poisson binomial test, Figure 7b) [34, 35], in concordance with previous studies of specific diseases [49, 50], and that these mutations affect a broad range of ligand-binding sites (Table S1).

Finally, we assess whether somatically mutated sites across human cancers overlap with putative binding positions across all human proteins as we might expect by random chance. Others have noted the propensity of cancer mutations to coincide with ligand interaction sites across smaller gene sets and in known driver genes in particular [13, 51]. We find that nearly a quarter of the 1,209 unique cancer-driving somatic mutations (DoCM, v3.2) [38], for instance, fall into confidently modeled putative ligand-binding sites inferred using InteracDome (Table S2), even though these sites constitute only ∼4.5% of the entire proteome and ∼12.8% of the proteome modeled by an interaction domain (*p* < 2.5e-20, binomial test). Moreover, when we repeat our global, site-based analysis, considering *all* somatically mutated sites across >10,000 tumor samples from 33 cancer types and using a much more comprehensive set of inferred binding sites than previous studies, we confirm the same trend. Sites harboring somatic missense mutations tend to coincide with inferred binding sites significantly *more* than expected by random chance (*p* < 6.9e-12, Poisson binomial test, Figure 7c), strongly suggesting that protein interaction perturbation is a frequent mechanism by which somatic mutations contribute to tumor fitness. Indeed, the somatic mutations affecting InteracDome-inferred putative binding sites have higher deleteriousness scores relative to non-binding mutations, as evaluated by various mutational impact predictors (*p* < 1e-7, Fisher’s exact tests, Table 1) [52]. However, unlike these other deleteriousness predictors, our InteracDome-inferred binding sites can not only be used to pinpoint potentially disease-relevant mutations, but can also be used to reason about their molecular, mechanistic impacts on protein interaction functionality.

**Table 1:** Fisher’s Exact Tests comparing deleteriousness predictions between binding and non-binding mutations. Each distinct somatic mutation in the pan-cancer dataset is classified as either binding (i.e., falls into an InteracDome-inferred, confidently modeled putative binding position in at least one human protein) or non-binding. Corresponding deleteriousness scores for each of these mutations were retrieved, where available, from the Database for Nonsynonymous SNPs’ Functional Predictions (v3.5) [52]; many mutations analyzed did not have corresponding deleteriousness scores for one or more predictors. Score thresholds to distinguish deleterious (“del.”) from tolerated (“tol.”) mutations were set as recommended by each method or to ≥0.5 when not specified for REVEL and MutPred scores. PolyPhen2 is abbreviated as “PP2”.

| Method | # Binding Muts. |  | # Non-Binding Muts. |  | Odds Ratio | *p*-value |
| --- | --- | --- | --- | --- | --- | --- |
|  | *del.* | *tol.* | *del.* | *tol.* |  |  |
| SIFT | 19,353 | 7,528 | 432,420 | 275,475 | 1.64 | 4e-297 |
| PP2, HDIV | 17,695 | 9,780 | 367,417 | 357,362 | 1.76 | ~0 |
| PP2, HVAR | 15,568 | 11,912 | 287,478 | 437,244 | 1.99 | ~0 |
| MutationTaster | 5,362 | 3,509 | 118,207 | 87,423 | 1.13 | 3e-8 |
| PROVEAN | 5,738 | 2,887 | 95,824 | 104,762 | 2.17 | 7e-259 |
| REVEL | 2,829 | 6,020 | 44,178 | 161,084 | 1.71 | 1e-109 |
| MutPred | 5,333 | 2,700 | 84,898 | 100,789 | 2.34 | 1e-291 |

The respective overlap (and lack thereof) of inferred binding sites with mutated or varying sites is significant even when considering only specific types of ligand binding in turn (Figure 7). Somatically mutated sites in particular appear to overlap with putative DNA-binding sites across proteins significantly more than expected, in accordance with what we know about impaired DNA repair functionality and perturbed regulatory processes in cancer [53]. Importantly, we continue to observe the same overall trends for sites exhibiting naturally occuring variation, Mendelian disease mutations, and somatic mutations when we consider alternate precision-based definitions of putative ligand-binding sites from the modelable-NR set (Figure S5). Overall, given that natural missense variants across healthy populations are depleted in putative binding sites, and Mendelian disease-associated missense mutations as well as somatic missense mutations across cancer tumors are enriched in them, computational approaches that leverage our InteracDome resource to identify perturbed interaction sites are likely to be highly relevant for identifying disease genes and informing disease mechanisms.

## DISCUSSION

We have introduced here a fully automated approach for aggregating protein co-complex structural data in the context of domains to reveal how positions within these domains are generally involved in mediating interactions with DNA, RNA, ions, peptides, metabolites, and other small molecules (Figure 1). This collection of 4,120 interaction domains, called the **InteracDome**, can be applied to pinpoint putative interaction sites in various proteins across species; here we show how a subset of 1,327 domains with confidently modeled ligand interactions can be used to infer functionally relevant interaction sites across the greatest proportion of human genes to date (Figure 6a).

Previously, interaction site information has been transferred from structurally modeled proteins to uncharacterized proteins with similar sequences or subsequences using various homology-based approaches [13, 46, 47]. Sequence motifs have also been semi-manually annotated with highly conserved metal ion binding or catalytic site information for use in identifying functional sites in new proteins, although such approaches are limited due to their rigid sequence match requirements [54]. Indeed, the ability of traditional homology-based approaches to infer binding information across larger, more diverse sets of protein sequences using existing structural templates is restricted in general because even as the number of resolved protein structures is increasing, the diversity of their sequences is not. Here, we develop a structurally-aware, domain-based approach to calculate real-valued binding propensities across individual domain positions. These probabilistically-modeled domain profiles are better able to capture conserved residues required for proper domain folding and thus can be used to accurately transfer binding site knowledge across a far more diverse set of proteins.

Determining the binding positions within these domains represents a challenging task due to biases inherent in structural data: structures often harbor confounding experimental artifacts, are dominated by non-cognate drug interactions, and can have highly redundant sequences to each other [24]. We show that by addressing each of these issues in turn, our systematic approach models binding positions across thousands of domains—including nearly all known DNA-, RNA-, and peptide-binding domains—that are highly indicative of ligand-binding positions in well characterized domains as well as in structural instances that were held out in cross-validation testing (Figure 2, Figure 5). Moreover, we find that aggregating domain co-complex structures to develop a general understanding of how a domain participates in interactions is superior to using only one or a few structures for this task (Figure 4b).

Substantial previous work has focused on detecting and characterizing the domain–domain interactions that mediate a number of protein–protein interactions across cellular interaction networks [55, 56, 19, 57, 58]. We make the distinction that in our work, we focus not on detecting the particular domain-mediated interfaces between specific protein partners, but rather on understanding which positions within protein domains mediate a variety of interactions in general with nucleic acids, peptides, metabolites, and a wide range of small molecules. Though we do not consider proteins in complex with other whole proteins here, our framework can be naturally extended to characterize domain–domain interactions in more depth.

Overall, we believe that our structural aggregation framework and resultant InteracDome resource lay the groundwork for many future domain-centric analyses and thus will be of broad use for the community. Not only can knowledge of domain binding positions be used to further group, subtype, subdivide, or functionally annotate domain families themselves, but the putative interaction sites inferred across proteins using the InteracDome should be relevant in understanding disease etiology. Indeed, we find that these sites are globally conserved across healthy human individuals yet preferentially perturbed in tumor samples and disease populations. As such, we anticipate that future approaches that utilize InteracDome to detect interaction-altering protein coding variants will be a great aid in both prioritizing disease-associated mutations as well as reasoning about their molecular effects.

## ACKNOWLEDGEMENTS

Thanks to members of the Singh lab for their insight and comments. Thanks especially to Dario Ghersi, Anton Periskov and Pawel Przytycki for helpful discussions regarding PDB files and TCGA mutation data. The results published here are in part based upon data generated by the TCGA Research Network: http://cancergenome.nih.gov/.

## SUPPORTING INFORMATION

**Figure S1.**
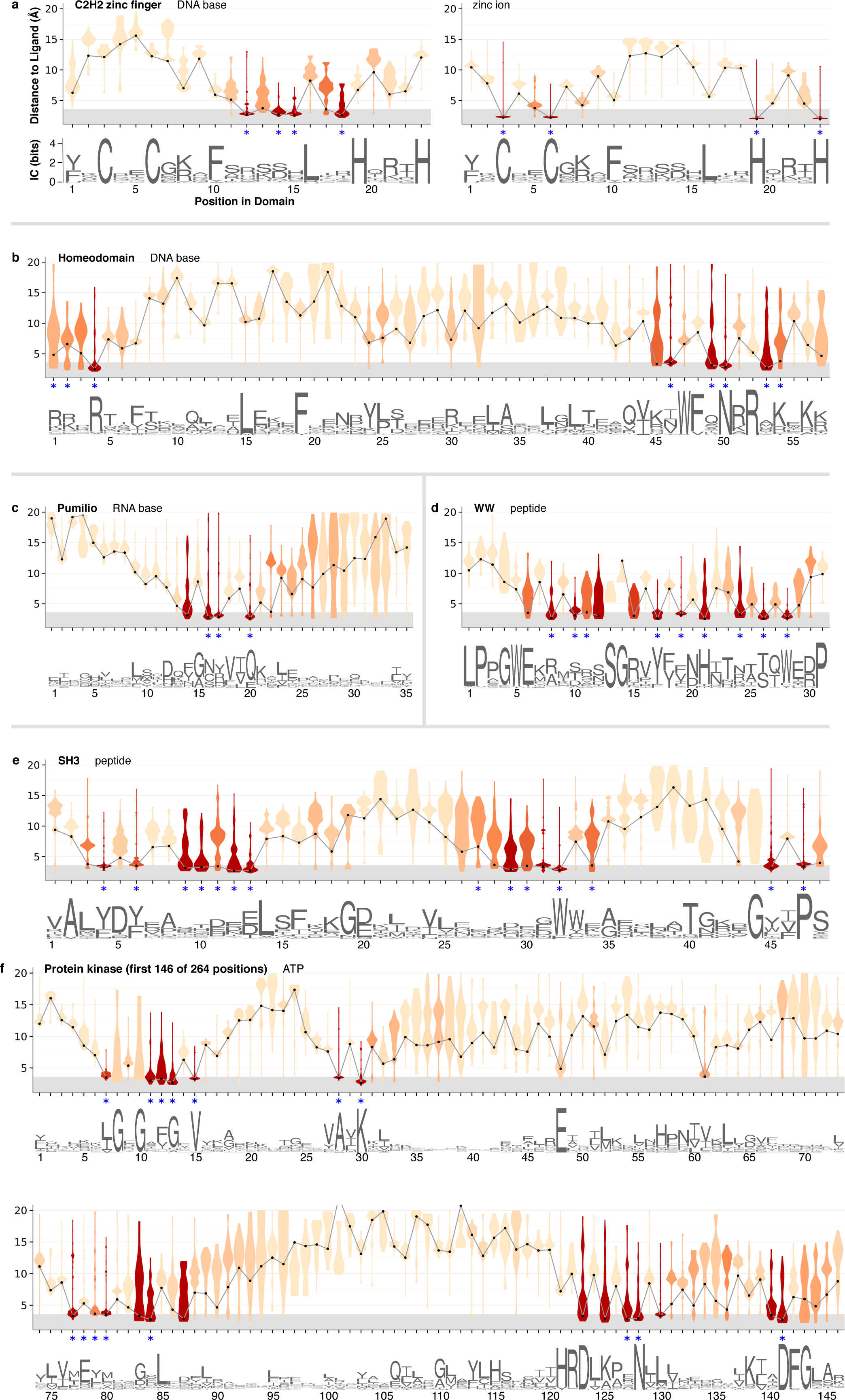
Examples of ligand-proximity scores across domains. For each position in each interaction domain, we show violin plots depicting the distribution of minimum receptor–ligand distances across instances in BioLiP, colored according to the fraction of the weighted distribution within 3.6Å. Gray lines connect the first deciles of each distribution. The *x*-axis is labeled with a sequence logo generated from the multiple sequence alignment of domain instances in BioLiP, where column height corresponds to information content. Previously identified ligand-contacting residues [59, 43, 26, 44, 45, 15] are marked with blue asterisks. The *x*-axis is labeled with a logo generated using Weblogo3 from the multiple sequence alignment of Pfam domain hits across BioLiP. The height of each column in the logos corresponds to the information content (IC) of that column; the logos in (a-f) are scaled equally according to the scale in (a). The particular interaction domains are: **(a)** Cys2-His2 zinc finger domain (PF00096), **(b)** Homeodomain (PF00046), **(c)** Pumilio domain (PF00806), **(d)** WW domain (PF00397), **(e)** SH3 domain (PF00018), and **(f)** the first 146 of 264 positions of protein kinase domain (PF00069).

**Figure S2.**
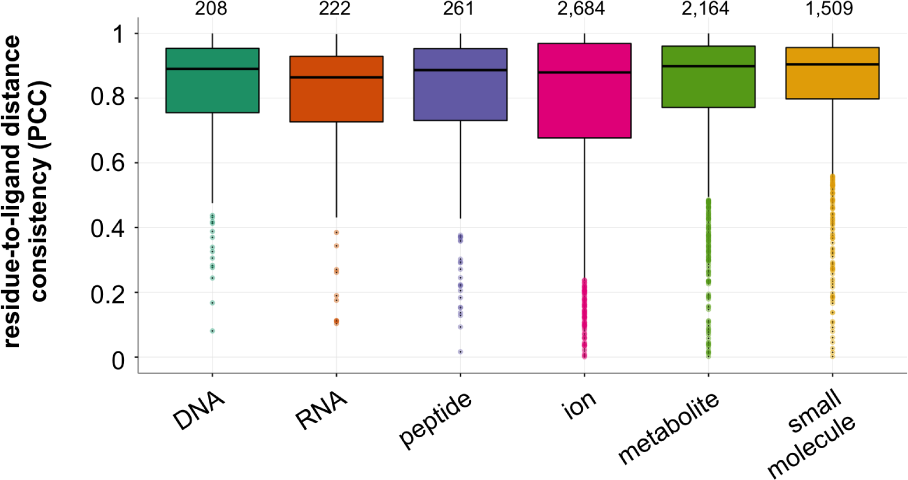
Domain-to-ligand distance consistencies between structural instances with < 90% sequence identity. Structural instances across BioLiP for each domain–ligand type are grouped by sequence similarity (≥ 90% identity), and then these groups are randomly split into two folds. Shown are Pearson’s correlation coefficients (PCCs) of the average residue-to-ligand distances across each domain position between the two folds. Total number of domain–ligand interactions included in each boxplot are listed above the corresponding distributions. The median PCC of domain–ligand interactions across these groups is 0.89.

**Figure S3.**
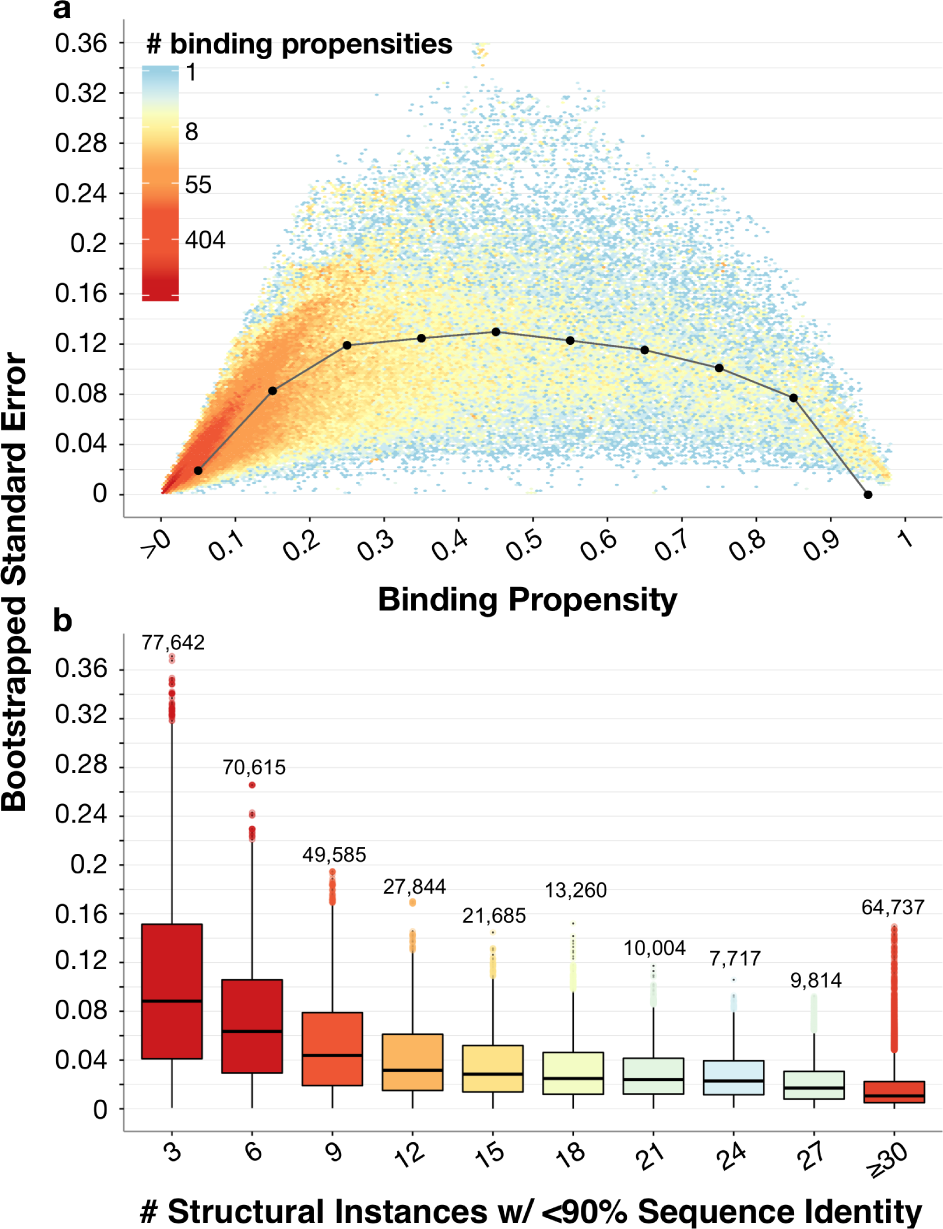
Standard errors of binding propensities obtained by bootstrapping groups of structural instances with ≥ 90% sequence identity. Structural instances across BioLiP for each domain–ligand type are grouped by sequence similarity (≥90% identity), and then these groups are randomly selected with replacement to generate 1,000 empirically bootstrapped sets of structural instances. **(a)** For each domain position with a positive binding propensity in each domain–ligand interaction pair, we plot its ligand-binding propensity (*x*-axis) and the standard error of this propensity (*y*-axis), computed as the standard deviation of its ligand-binding propensity as measured over 1,000 bootstrap samples. Distribution medians at each binding propensity decile are shown as black dots and are connected by gray lines for visual effect. **(b)** Bootstrapped standard errors decrease as the number of domain–ligand structural instances with <90% sequence identity increase. Boxplots are colored according to the relative size of each distribution; the number of total domain positions, across domain–ligand type pairs, is listed above each boxplot.

**Figure S4.**
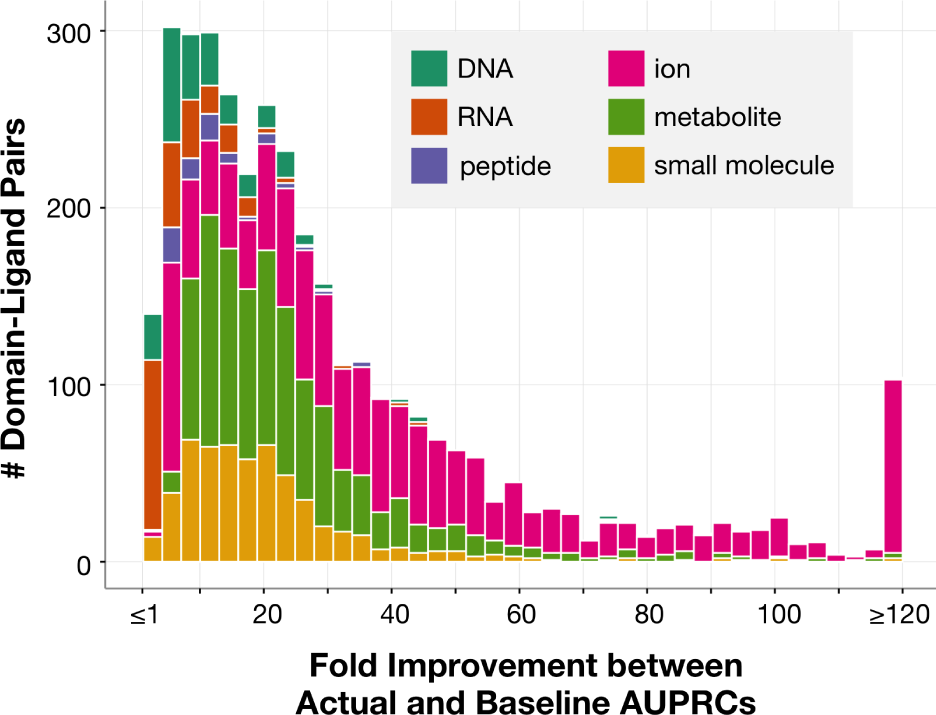
Cross-validation testing of binding propensities where structural instances in distinct folds have < 90% sequence identity to each other. Structural instances across BioLiP for each domain–ligand type are grouped by sequence similarity (≥90% identity), and then these groups are randomly split into up to 10 folds. Accuracy of each domain–ligand interaction with 2+ groups of structural instances is measured as the average area under the precision-recall curve (AUPRC) in cross-validation. For each domain–ligand pair, we compute the fold change between the actual AUPRC and a baseline AUPRC corresponding to the fraction of binding positions in held-out sets.

**Figure S5.**
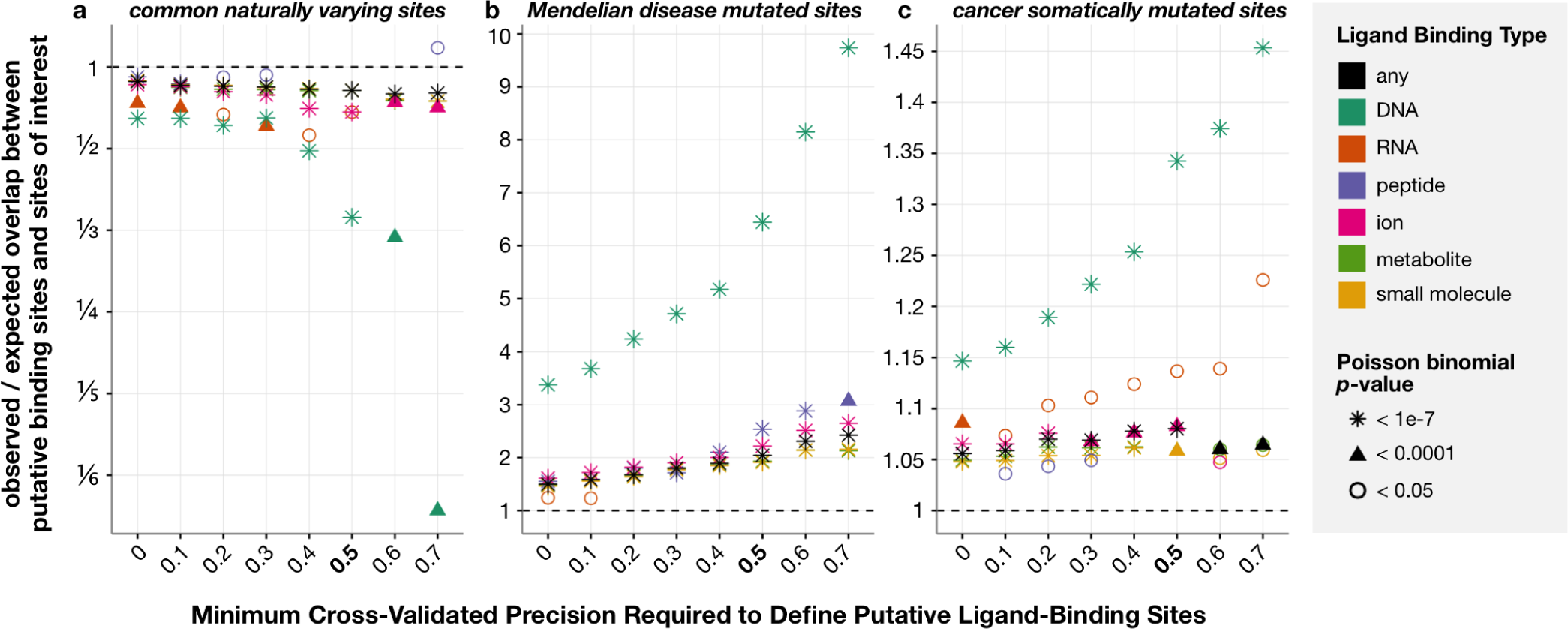
Natural variants show opposite trends from disease mutations to overlap with putative ligand-binding sites. Putative ligand binding sites are inferred across human proteins using domains from the modelable-NR set. Specifically, for precision thresholds between 0 and 0.7 (*x*-axis), protein positions overlapping domain match states whose binding propensities resulted in at least that precision in cross-validation testing (see Materials and Methods) are considered to be putative binding sites. We compute the significance of the overlap between InteracDome-inferred binding sites and other sites of interest using the Poisson binomial distribution. Bolded along the *x*-axis is the value used to define putative ligand-binding sites in the main text (i.e., confidently modeled interactions); shown along the *y*-axis is the fold change between the observed number of overlapping sites (K) and the expected number of overlapping sites (E[K]). **(a)** Putative ligand-binding sites exhibit a significant lack of overlap with commonly varying sites across human proteins. Each point corresponds to the fold change between these values for a particular type of ligand-binding site (indicated by its color) for a particular precision-based definition of putative binding site. The shape of each point corresponds to its computed *p*-value. Fold change values that have corresponding *p*-values ≥0.05 are not shown. **(b)** Conversely, putative ligand-binding sites across human proteins overlap significantly with sites harboring Mendelian disease mutations; points are colored and shaped as in (a). **(c)** Protein sites harboring a missense cancer somatic mutation also overlap significantly with putative ligand-binding sites.

**Table S1.** Counts of Mendelian disease mutations affecting particular types of ligand-binding sites. We consider 30,154 distinct Mendelian-associated missense mutations across 26,434 protein sites in 2,749 proteins. We define putative ligand-binding sites in two ways. First, for each domain–ligand type pair in the modelable-NR set, we find matches to the domain in canonical human protein isoform sequences, and consider any protein residue that overlaps with a domain match state whose binding propensity is positive to be a putative ligand-binding site (i.e., modelable-NR interactions); 12% of mutations affect these sites. Second, we consider any protein residue that overlaps with a domain match state whose binding propensity resulted in a precision of at least 0.5 in cross-validation testing to be a putative ligand-binding site (i.e., confidently modeled interactions, as in the main text); 4% of mutations affect these sites. Columns (from left to right) are the type of ligand interaction, total mutations to affect modelable-NR binding sites as described here, total mutations to affect confidently modeled binding sites, total number of mutated confidently modeled binding sites, and total number of proteins with mutated confidently modeled binding sites. Note that the sets of putative nucleic acid base and backbone binding sites are overlapping.

| Binding Site Type | Modelable-NR Interactions | Confidently Modeled Interactions |  |  |
| --- | --- | --- | --- | --- |
|  | *Total Mutations* | *Total Mutations* | *Affected Sites* | *Affected Proteins* |
| *any* | 3,718 | 1,152 | 964 | 415 |
| *DNA* | 284 | 118 | 94 | 48 |
| *DNA base* | 157 | 81 | 68 | 40 |
| *DNA backbone* | 284 | 116 | 92 | 45 |
| *RNA* | 70 | 15 | 14 | 6 |
| *RNA base* | 40 | 3 | 3 | 3 |
| *RNA backbone* | 70 | 17 | 15 | 6 |
| *peptide* | 736 | 74 | 68 | 46 |
| *ion* | 974 | 273 | 218 | 107 |
| *metabolite* | 2,466 | 673 | 565 | 266 |
| *small molecule* | 2,880 | 743 | 625 | 291 |

**Table S2.**
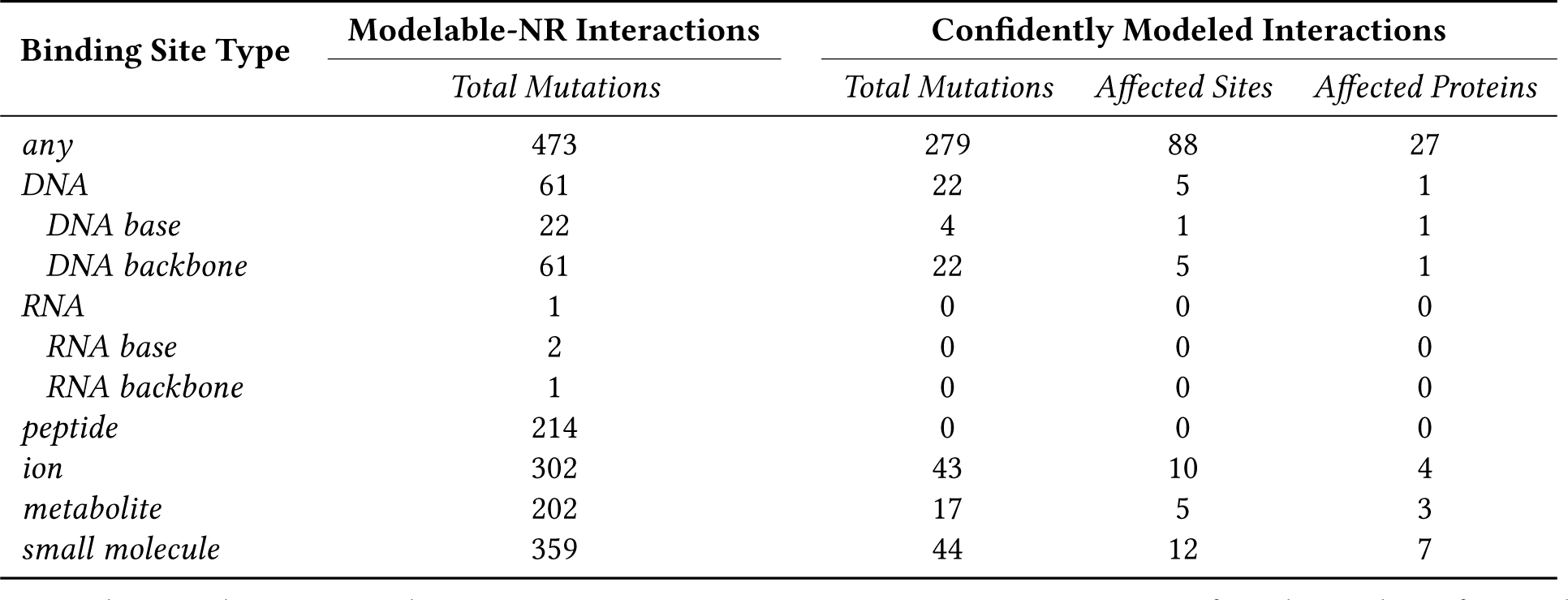
Counts of known cancer-driving mutations affecting particular types of ligand-binding sites. We consider 1,209 distinct cancer driver missense mutations across 571 protein sites in 128 proteins from the Database of Curated Mutations. We define putative ligand-binding sites in two ways. First, for each domain–ligand type pair in the modelable-NR set, we find matches to the domain in the longest human protein isoform sequences, and consider any protein residue that overlaps with a domain match state whose binding propensity is positive to be a putative ligand-binding site (i.e., modelable-NR interactions); 39% of mutations affect these sites. Second, we consider any protein residue that overlaps with a domain match state whose binding propensity resulted in a precision of at least 0.5 in cross-validation testing to be a putative ligand-binding site (i.e., confidently modeled interactions, as in the main text); 23% of mutations affect these sites. Columns (from left to right) are the type of ligand interaction, total mutations to affect modelable-NR binding sites as described here, total mutations to affect confidently modeled binding sites, total number of mutated confidently modeled binding sites, and total number of proteins with mutated confidently modeled binding sites. Note that the sets of putative nucleic acid base and backbone binding sites are overlapping.

## References

[1] Finn, R.D., Clements, J., and Eddy, S.R. (2011) HMMER web server: Interactive sequence similarity searching. Nucleic Acids Res, 39, W29–W37.

[2] Finn, R.D., Bateman, A., Clements, J., Coggill, P., Eberhardt, R.Y., Eddy, S.R., Heger, A., Hetherington, K., Holm, L., Mistry, J., Sonnhammer, E.L.L., Tate, J., and Punta, M. (2014) Pfam: the protein families database. Nucleic Acids Res, 42, D222–D230.

[3] Letunic, I. and Bork, P. (2018) 20 years of the SMART protein domain annotation resource. Nucleic Acids Res, 46, D493–D496.

[4] Marchler-Bauer, A., Bo, Y., Han, L., He, J., Lanczycki, C.J., Lu, S., Chitsaz, F., Derbyshire, M.K., Geer, R.C., Gonzales, N.R., et al. (2017) CDD/SPARCLE: functional classification of proteins via subfamily domain architectures. Nucleic Acids Res, 45, D200–D203.

[5] Ochoa, A., Llinas, M., and Singh, M. (2011) Using context to improve protein domain identification. BMC Bioinformatics, 12, 90.

[6] Vaquerizas, J.M., Kummerfeld, S.K., Teichmann, S.A., and Luscombe, N.M. (2009) A census of human transcription factors: Function, expression and evolution. Nat Rev Genet, 10, 252–263.

[7] Gerstberger, S., Hafner, M., and Tuschl, T. (2014) A census of human RNA-binding proteins. Nat Rev Genet, 15, 829–845.

[8] Cohen, G., Ren, R., and Baltimore, D. (1995) Modular binding domains in signal transduction proteins. Cell, 80, 237–248.

[9] Forslund, K. and Sonnhammer, E.L. (2008) Predicting protein function from domain content. Bioinformatics, 24, 1681–1687.

[10] Kim, P.M., Lu, L.J., Xia, Y., and Gerstein, M.B. (2006) Relating three-dimensional structures to protein networks provides evolutionary insights. Science, 314, 1938–1941.

[11] Betts, M.J., Lu, Q., Jiang, Y., Drusko, A., Wichmann, O., Utz, M., Valtierra-Gutiérrez, I.A., Schlesner, M., Jaeger, N., Jones D.T., et al. (2015) Mechismo: predicting the mechanistic impact of mutations and modifications on molecular interactions. Nucleic Acids Res, 43, e10.

[12] Hosur, R., Xu, J., Bienkowska, J., and Berger, B. (2011) iWRAP: An interface threading approach with application to prediction of cancer-related protein-protein interactions. J Mol Biol, 405, 1295–1310.

[13] Ghersi, D. and Singh, M. (2014) Interaction-based discovery of functionally important genes in cancers. Nucleic Acids Res, 42, e18.

[14] Winter, C., Henschel, A., Tuukkanen, A., and Schroeder, M. (2012) Protein interactions in 3D: from interface evolution to drug discovery. J Struct Biol, 179, 347–358.

[15] Hanks, S.K. and Hunter, T. (1995) Protein kinases 6. The eukaryotic protein kinase superfamily: kinase (catalytic) domain structure and classification. The FASEB Journal, 9, 576–596.

[16] Persikov, A.V., Osada, R., and Singh, M. (2009) Predicting DNA recognition by Cys2His2 zinc finger proteins. Bioinformatics, 25, 22–29.

[17] Pabo, C.O., Peisach, E., and Grant, R.A. (2001) Design and selection of novel Cys2His2 zinc finger proteins. Ann Rev Biochem, 70, 313–340.

[18] Barrera, L.A., Vedenko, A., Kurland, J.V., Rogers, J.M., Gisselbrecht, S.S., Rossin, E.J., Woodard, J., Mariani, L., Kock, K.H., Inukai, S., et al. (2016) Survey of variation in human transcription factors reveals prevalent DNA binding changes. Science, 351, 1450–1454.

[19] Mosca, R., Ceol, A., Stein, A., Olivella, R., and Aloy, P. (2014) 3did: a catalog of domain-based interactions of known three-dimensional structure. Nucleic Acids Res, 42, D374–D379.

[20] Finn, R.D., Miller, B.L., Clements, J., and Bateman, A. (2014) iPfam: a database of protein family and domain interactions found in the Protein Data Bank. Nucleic Acids Res, 42, D364–D373.

[21] Isserlin, R., El-Badrawi, R.A., and Bader, G.D. (2011) The Biomolecular Interaction Network Database in PSI-MI 2.5. Database, 2011, baq037.

[22] Xu, Q. and Dunbrack, R.L. (2012) Assignment of protein sequences to existing domain and family classification systems: Pfam and the PDB. Bioinformatics, 28, 2763–2772.

[23] Bashton, M., Nobeli, I., and Thornton, J.M. (2008) PROCOGNATE: a cognate ligand domain mapping for enzymes. Nucleic Acids Res, 36, D618–D622.

[24] Yang, J., Roy, A., and Zhang, Y. (2013) BioLiP: a semi-manually curated database for biologically relevant ligand–protein interactions. Nucleic Acids Res, 41, D1096–D1103.

[25] Eddy, S.R. (2011) Accelerated profile HMM searches. PLoS Comput Biol, 7, e1002195.

[26] Wang, X., McLachlan, J., Zamore, P.D., and Hall, T.M.T. (2002) Modular recognition of RNA by a human pumilio-homology domain. Cell, 110, 501–512.

[27] Rogers, D.J.and Tanimoto, T.T. (1960) A computer program for classifying plants. Science, 132, 1115–1118.

[28] O’Boyle, N.M., Banck, M., James, C.A., Morley, C., Vandermeersch, T., and Hutchison, G.R. (2011) Open Babel: an open chemical toolbox. J Cheminform, 3, 33.

[29] Wishart, D.S., Jewison, T., Guo, A.C., Wilson, M., Knox, C., Liu, Y., Djoumbou, Y., Mandal, R., Aziat, F., Dong, E., et al. (2013) HMDB 3.0–the Human Metabolome DataBase in 2013. Nucleic Acids Res, 41, D801–D807.

[30] Wishart, D.S., Knox, C., Guo, A.C., Shrivastava, S., Hassanali, M., Stothard, P., Chang, Z., and Woolsey, J. (2006) DrugBank: a comprehensive resource for in silico drug discovery and exploration. Nucleic Acids Res, 34, D668–D672.

[31] Henikoff, S. and Henikoff, J.G. (1994) Position-based sequence weights. J Mol Biol, 243, 574–578.

[32] Persikov, A.V. and Singh, M. (2011) An expanded binding model for Cys2 His2 zinc finger protein–DNA interfaces. Phys Biol, 8, e035010.

[33] Lek, M., Karczewski, K.J., Minikel, E.V., Samocha, K.E., Banks, E., Fennell, T., O’Donnell-Luria, A.H., Ware, J.S., Hill, A.J., Cummings, B.B., et al. (2016) Analysis of protein-coding genetic variation in 60,706 humans. Nature, 536, 285–291.

[34] The UniProt Consortium (2012) Reorganizing the protein space at the Universal Protein Resource (UniProt). Nucleic Acids Res, 40, D71–D75.

[35] Martin, A.C.R. Mapping OMIM mutations to SwissProt D.J. (2011; accessed Apr. 24, 2017).

[36] Fan, Y., Xi, L., Hughes, D.S.T., Zhang, J., Zhang, J., Futreal, P.A., Wheeler, D.A., and Wang, W. (2016) MuSE: accounting for tumor heterogeneity using a sample-specific error model improves sensitivity and specificity in mutation calling from sequencing data. Genome Biol, 17, 178.

[37] Grossman, R.L., Heath, A.P., Ferretti, V., Varmus, H.E., Lowy, D.R., Kibbe, W.A., and Staudt, L.M. (2016) Toward a shared vision for cancer genomic data. N Engl J Med, 375, 1109–1112.

[38] Ainscough, B.J., Griffith, M., Coffman, A.C., Wagner, A.H., Kunisaki, J., Choudhary, M.N., McMichael, J.F., Fulton, R.S., Wilson, R.K., Griffith, O.L., et al. (2016) DoCM: a Database of Curated Mutations in cancer. Nat Meth, 13, 806–807.

[39] Hong, Y. (2013) On computing the distribution function for the Poisson binomial distribution. Comput Stat Data Anal, 59, 41–51.

[40] Luscombe, M.N., Austin, E.S., Berman, M.H., and Thornton, M.J. (2000) An overview of the structures of protein-DNA complexes. Genome Biol, 1, previews001.1.

[41] Lunde, B.M., Moore, C., and Varani, G (2007) RNA-binding proteins: modular design for efficient function. Nat Rev Mol Cell Biol, 8, 479–490.

[42] Sudha, G., Singh, P., Swapna, L.S., and Srinivasan, N. (2015) Weak conservation of structural features in the interfaces of homologous transient protein–protein complexes. Protein Sci, 24, 1856–1873.

[43] Noyes, M.B., Christensen, R.G., Wakabayashi, A., Stormo, G.D., Brodsky, M.H., and Wolfe, S.A. (2008) Analysis of homeodomain specificities allows the family-wide prediction of preferred recognition sites. Cell, 133, 1277–1289.

[44] Kato, Y., Ito, M., Kawai, K., Nagata, K., and Tanokura, M. (2002) Determinants of ligand specificity in groups I and IV WW domains as studied by surface plasmon resonance and model building. J Biol Chem, 277, 10173–10177.

[45] Saksela, K. and Permi, P. (2012) SH3 domain ligand binding: What’s the consensus and where’s the specificity?. FEBS Lett, 586, 2609–2614.

[46] Gress, A., Ramensky, V., Büch, J., Keller, A., and Kalinina, O.V. (2016) StructMAn: annotation of single-nucleotide polymorphisms in the structural context. Nucleic Acids Res, 44, W463–W468.

[47] Pieper, U., Webb, B.M., Dong, G.Q., Schneidman-Duhovny, D., Fan, H., Kim, S.J., Khuri, N., Spill, Y.G., Weinkam, P., Hammel, M., Tainer, J.A., Nilges, M., and Sali, A. (2014) ModBase, a database of annotated comparative protein structure models and associated resources. Nucleic Acids Res, 42, D336–D346.

[48] Berman, H.M., Westbrook, J., Feng, Z., Gilliland, G., Bhat, T.N., Weissig, H., Shindyalov, I.N., and Bourne, P.E. (2000) The Protein Data Bank. Nucleic Acids Res, 28, 235–242.

[49] Sahni, N., Yi, S., Zhong, Q., Jailkhani, N., Charloteaux, B., Cusick, M.E., and Vidal, M. (2013) Edgotype: A fundamental link between genotype and phenotype. Curr Opin Genet Dev, 23, 649–657.

[50] Sahni, N., Yi, S., Taipale, M., Bass, J.I.F., Coulombe-Huntington, J., Yang, F., Peng, J., Weile, J., Karras, G.I., Wang, Y., et al. (2015) Widespread macromolecular interaction perturbations in human genetic disorders. Cell, 161, 647–660.

[51] Gress, A., Ramensky, V.E., and Kalinina, O.V. (2017) Spatial distribution of disease-associated variants in three-dimensional structures of protein complexes. Oncogenesis, 6, e380.

[52] Liu, X., Wu, C., Li, C., and Boerwinkle, E. (2016) dbNSFP v3.0: A one-stop database of functional predictions and annotations for human nonsynonymous and splice-site SNVs. Hum Mutat, 37, 235–241.

[53] Jeggo, P.A., Pearl, L.H., and Carr, A.M. (2016) DNA repair, genome stability and cancer: a historical perspective. Nat Rev Cancer, 16, 35–42.

[54] Sigrist, C.J.A., De Castro, E., Langendijk-Genevaux, P.S., Le Saux, V., Bairoch, A., and Hulo, N. (2005) ProRule: a new database containing functional and structural information on PROSITE profiles. Bioinformatics, 21, 4060–4066.

[55] Shoemaker, B.A., Panchenko, A.R., and Bryant, S.H. (2006) Finding biologically relevant protein domain interactions: conserved binding mode analysis. Protein Sci, 15(2), 352–361.

[56] Ooi, H.S., Schneider, G., Chan, Y.L., Lim, T.T., Eisenhaber, B., and Eisenhaber, F. (2010) Databases of protein-protein interactions and complexes. Methods Mol Biol, 609, 145–159.

[57] Raghavachari, B., Tasneem, A., Przytycka, T.M., and Jothi, R. (2008) DOMINE: a database of protein domain interactions. Nucleic Acids Res, 36, D656–661.

[58] Mosca, R., Ceol, A., and Aloy, P. (2013) Interactome3D: adding structural details to protein networks. Nat Methods, 10, 47–53.

[59] Wolfe, S.A., Nekludova, L., and Pabo, C.O. (2000) DNA recognition by Cys2His2 zinc finger proteins. Ann Rev Bioph Biom, 29, 183–212.

